## Supplementary figures and methods for "Immunosuppression in the prostate tumor microenvironment is tied to androgen deprivation therapy-resistant club-like cells"

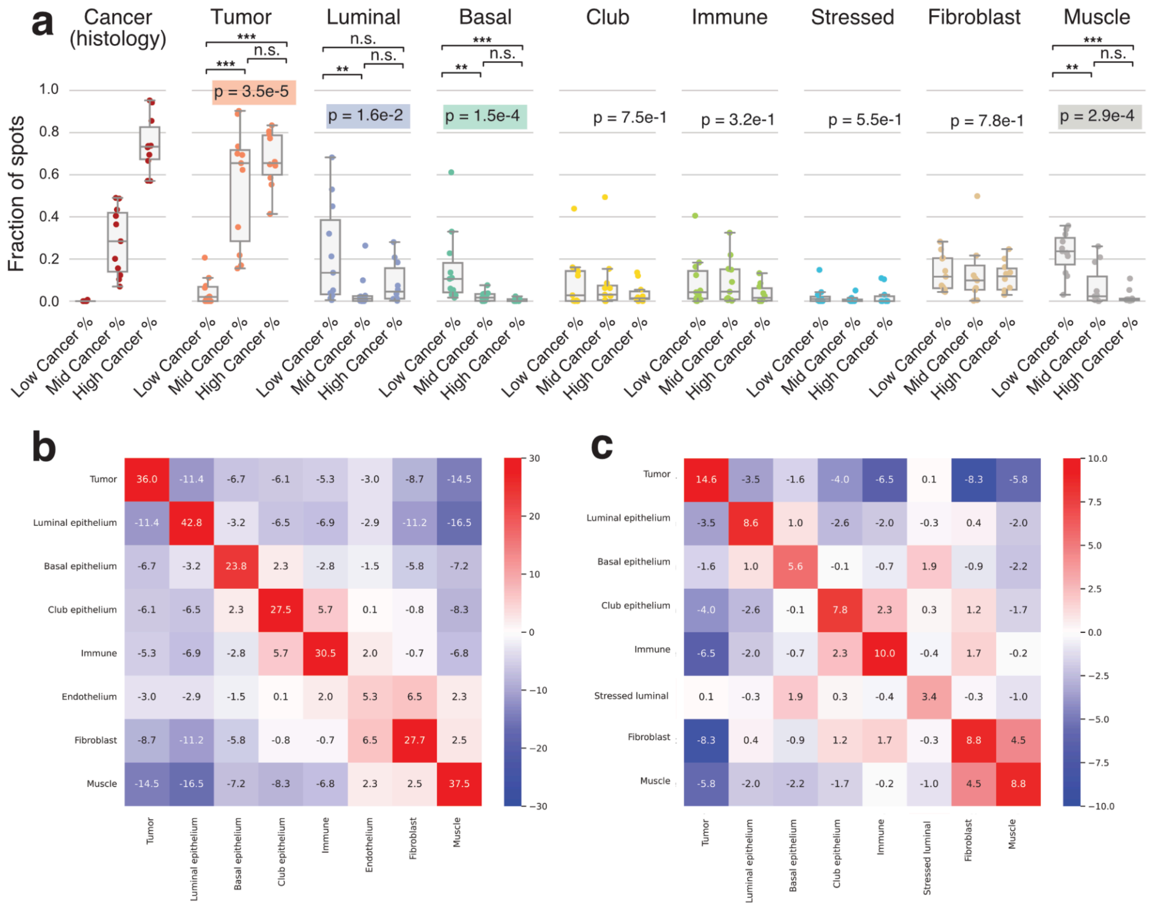


**Supplementary Figure S1. a)** Region fraction distributions divided by the percentage of ISUP-grade tumor spots. Kruskal-Wallis test p-value is displayed. Asterisks indicate Wilcoxon rank-sum test p-values; *: $p < 0.05$ ; **: $p < 0.01$ ; ***: $p < 0.001$. Mean region proximity scores across the **b)** the discovery and **c)** the validation cohorts. A higher z-score indicates a larger likelihood of the regions localizing in adjacent spots. Low: cancer % (% of spots labeled as ISUP1-5, ISUPX, ISUPY or PNI) ≤ 0.05; Mid: 0.05 < cancer % ≤ 0.50; High: 0.50 < cancer %.

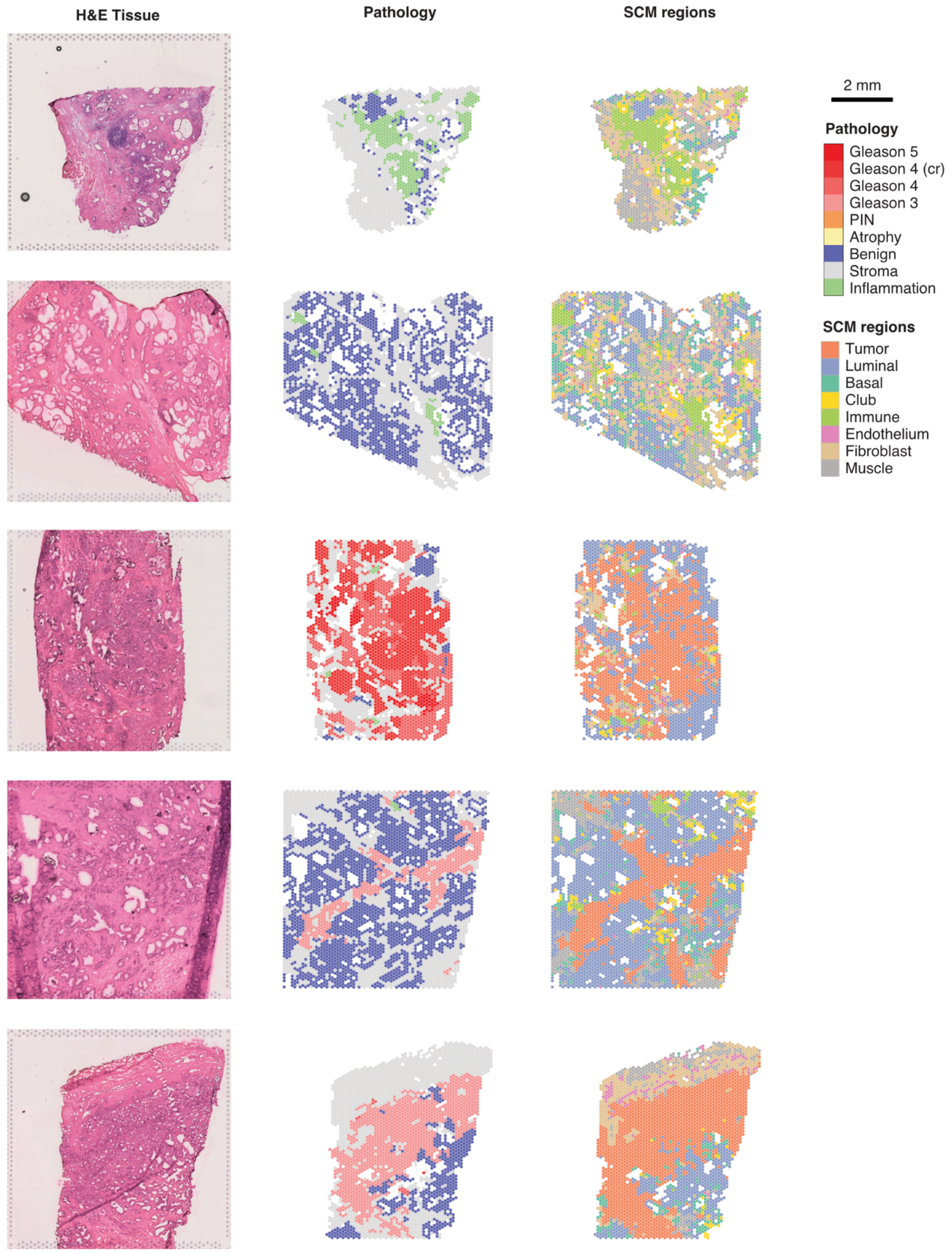


**Supplementary Figure S2**. Side-by-side H&E staining, histopathology assessment, and SCM regions for a representative set of discovery cohort samples. The first two samples are BPH, samples three and four are untreated tumors, and sample five is a bicalutamide-treated tumor.

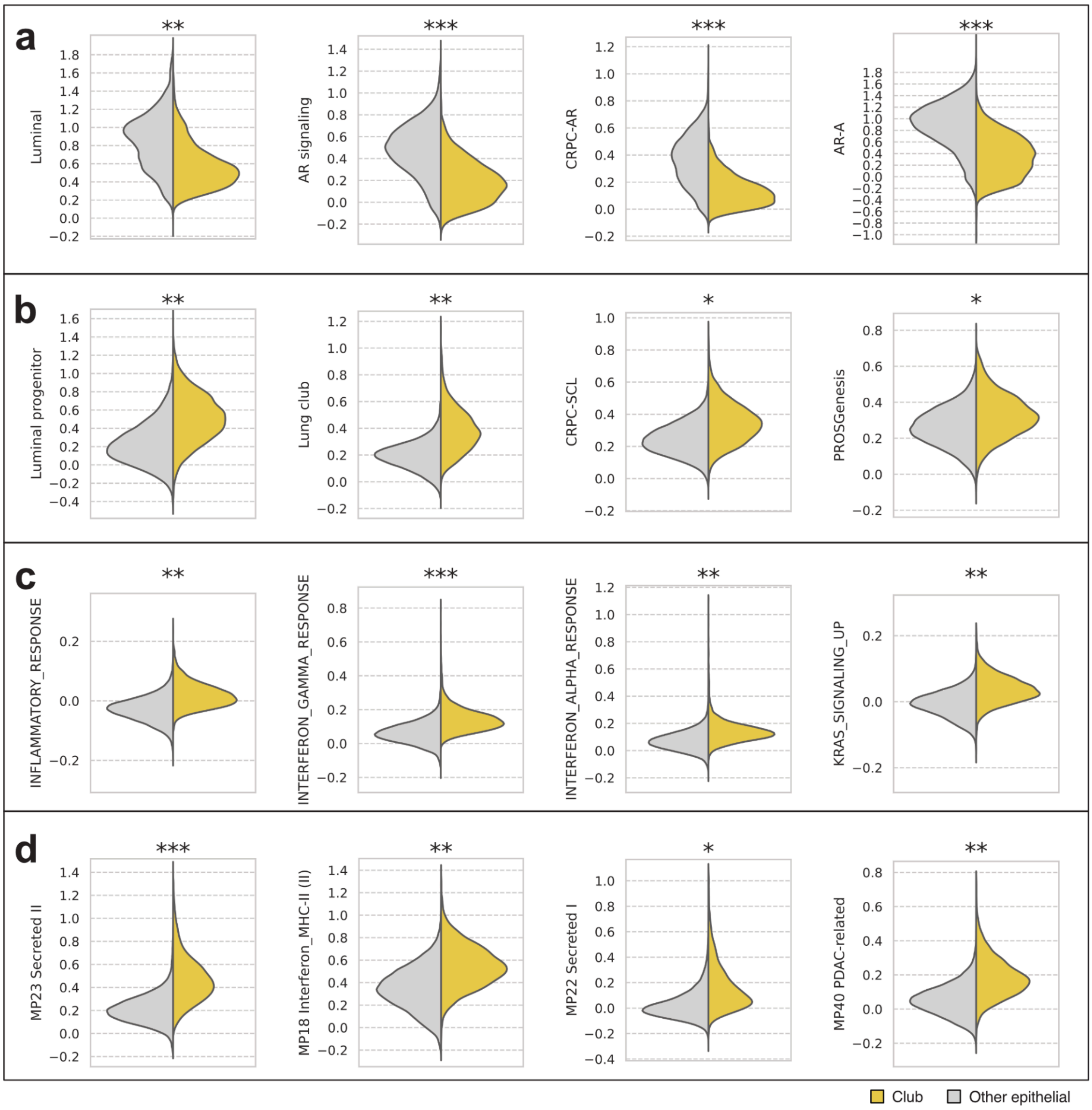


**Supplementary Figure S3.** Spotwise gene set activity scores in the Club region compared to other epithelial regions. **a)** Androgen signaling activity-related gene sets. **b)** Club-, stem-cell-, and progenitor-like gene sets. **c)** Inflammation-related MSigDB hallmark gene sets. **d)** Pan-cancer meta-program[1] gene sets. Asterisks indicate two-sided independent samples t-test p-value ($p < 0.05$) and effect size: *: Other 70th percentile < Club region mean; **: Other 80th percentile < Club region mean; ***: Other 90th percentile < Club region mean.

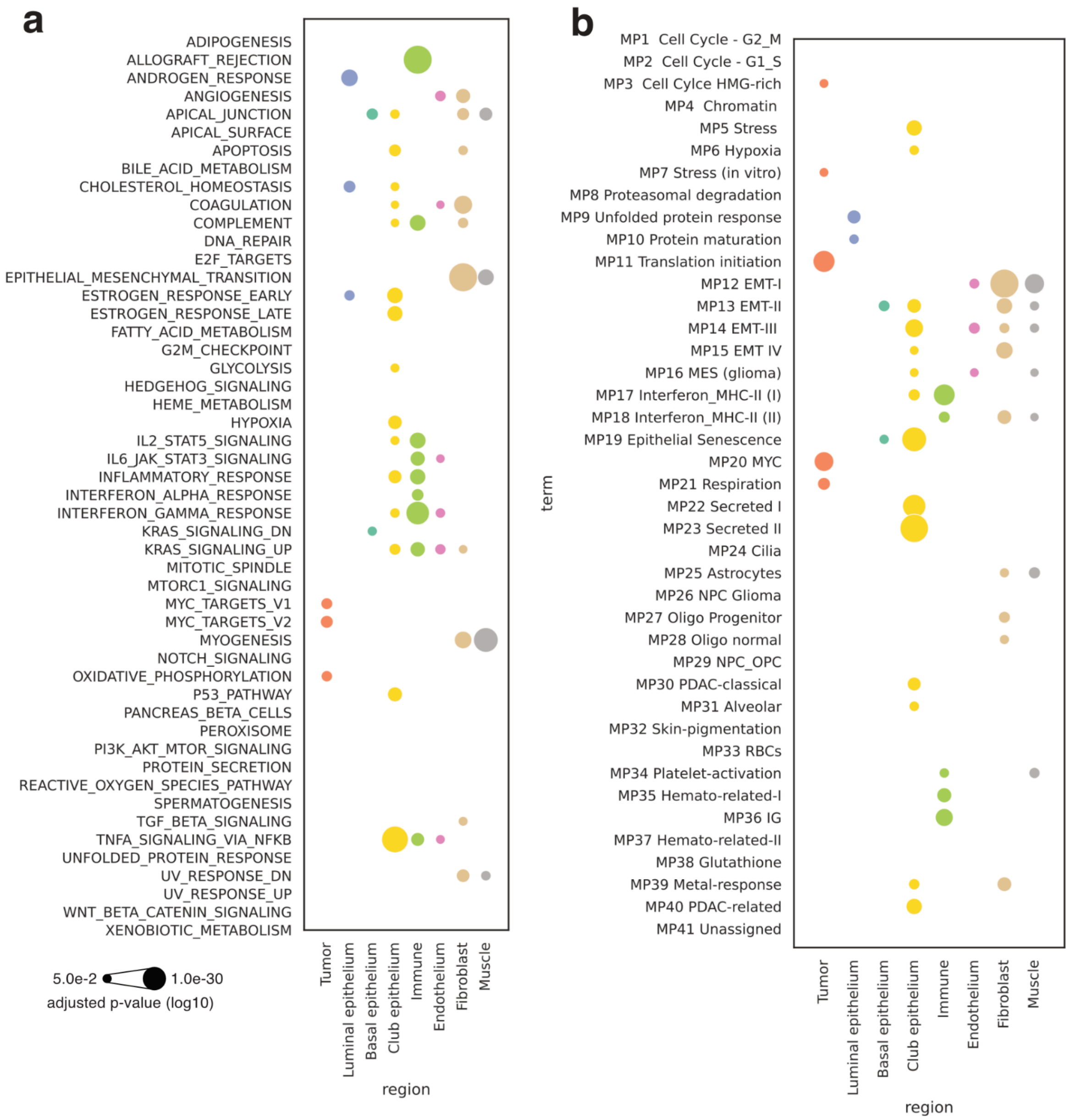


**Supplementary Figure S4.** Region markers enrichment analysis results for **a)** MSigDB Hallmark **b)** and pan-cancer meta-program gene sets. Dot size represents the Benjamini-Hochberg adjusted p-value of a one-sided Fisher's exact test. Colors represent regions.

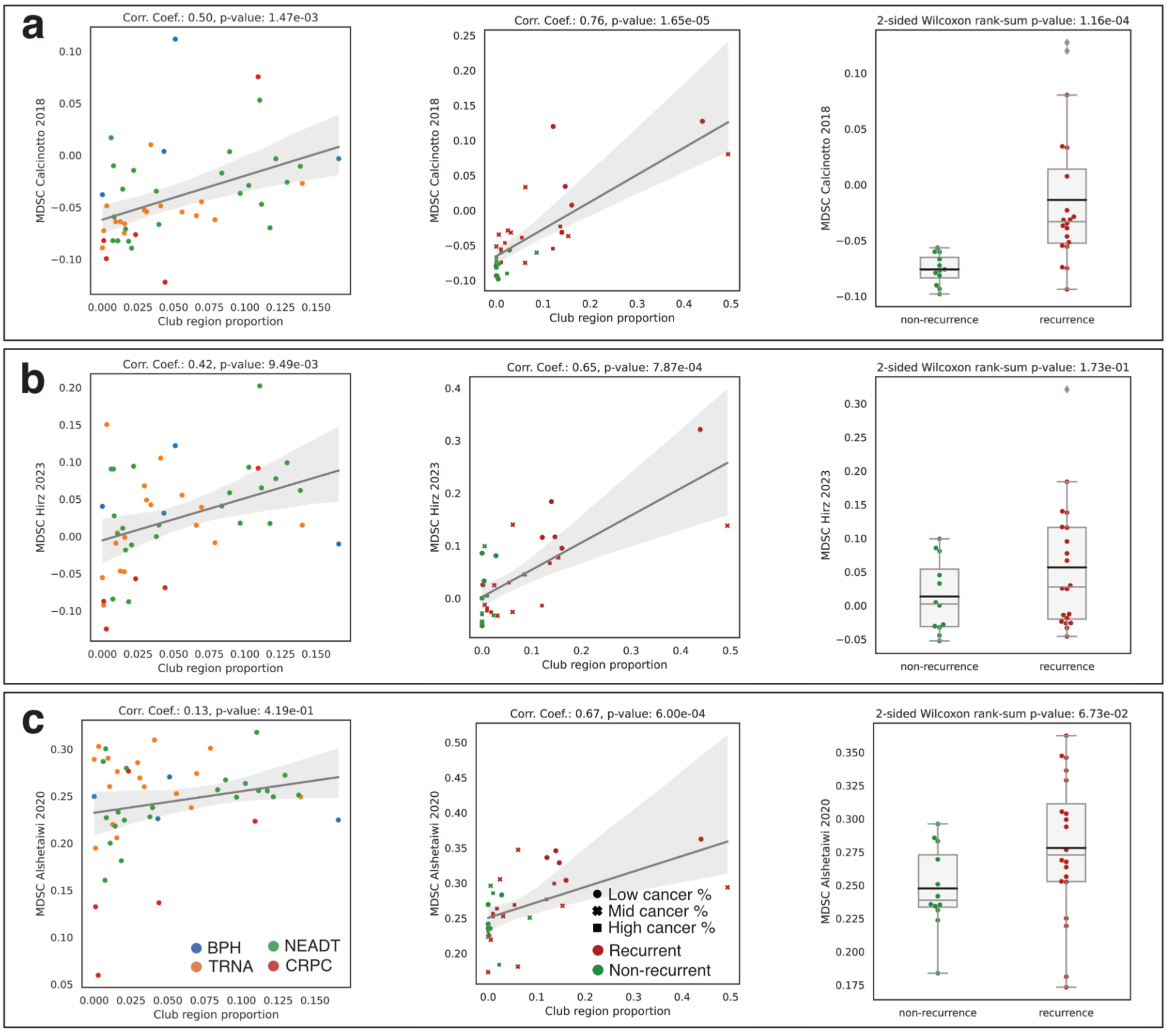


**Supplementary Figure S5.** Correlation between the proportion of Club region spots and the mean gene set activity score in the non-Club region spots for discovery (right) and the validation cohorts (middle). Spearman correlation coefficient and Benjamini-Hochberg corrected p-values are displayed. The boxplots depict score distributions of non-recurrent and recurrent samples and two-sided Wilcoxon rank-sum test p-values. **a**) Calcinotto et al. 2018 MDSC signature[2] **b)** Hirz et al. 2023 MDSC signature[3] **c)** Alshetaiwi et al. 2020 MDSC signature[4]

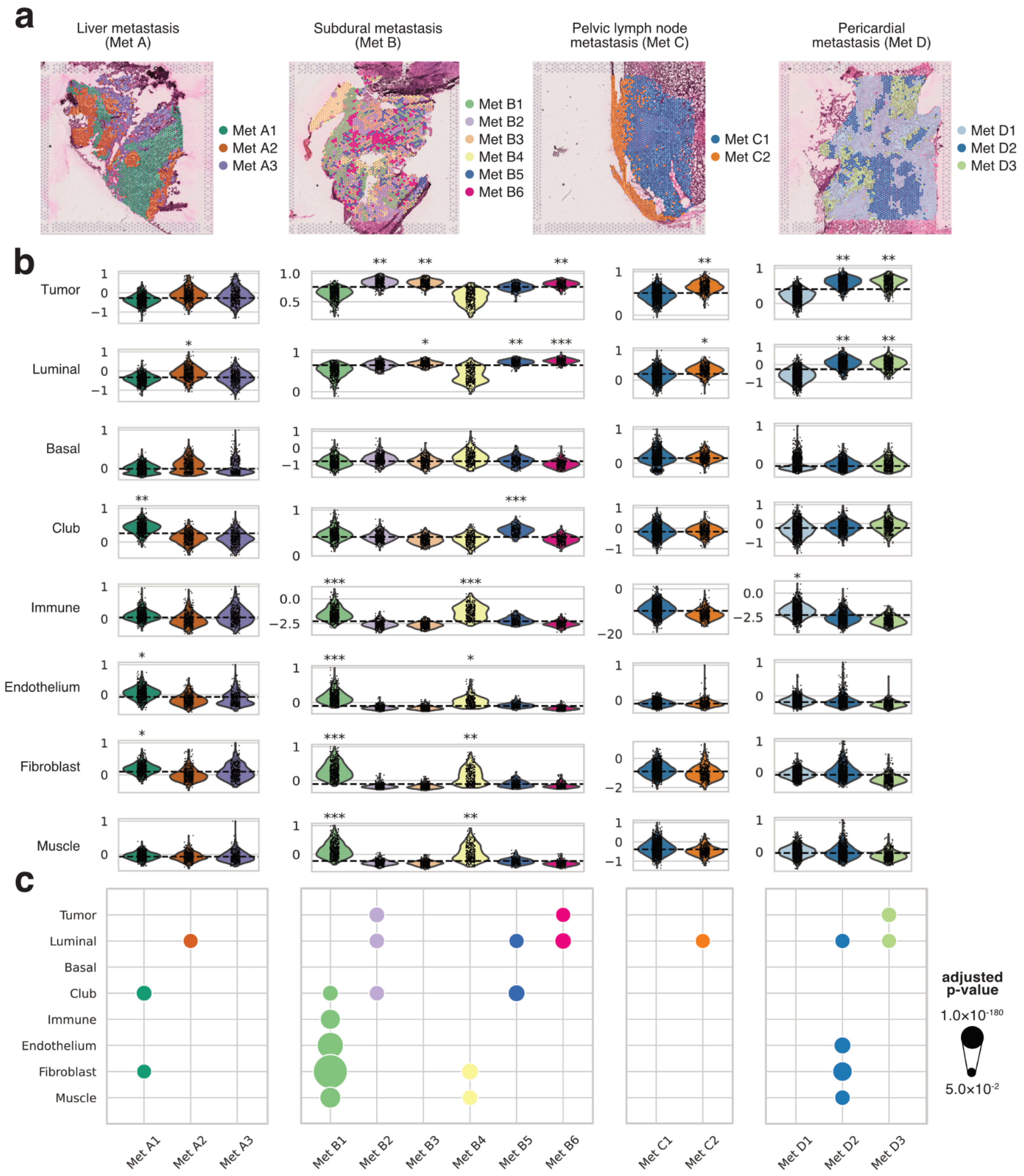


**Supplementary Figure S6.** Spatial transcriptomics data from metastatic prostate cancer samples. **a)** Expression-based clustering overlaid on spatial sections. **b)** Cluster-specific gene set scores. The dashed line marks the overall score median. Asterisks indicate two-sided independent samples t-test p-value ($p < 0.05$) and effect size (*: cluster 30th percentile > overall median; **: cluster 20th percentile > overall median; *: cluster 10th percentile > overall median). **c)** Adjusted overrepresentation p-values among each cluster's overexpressed genes.

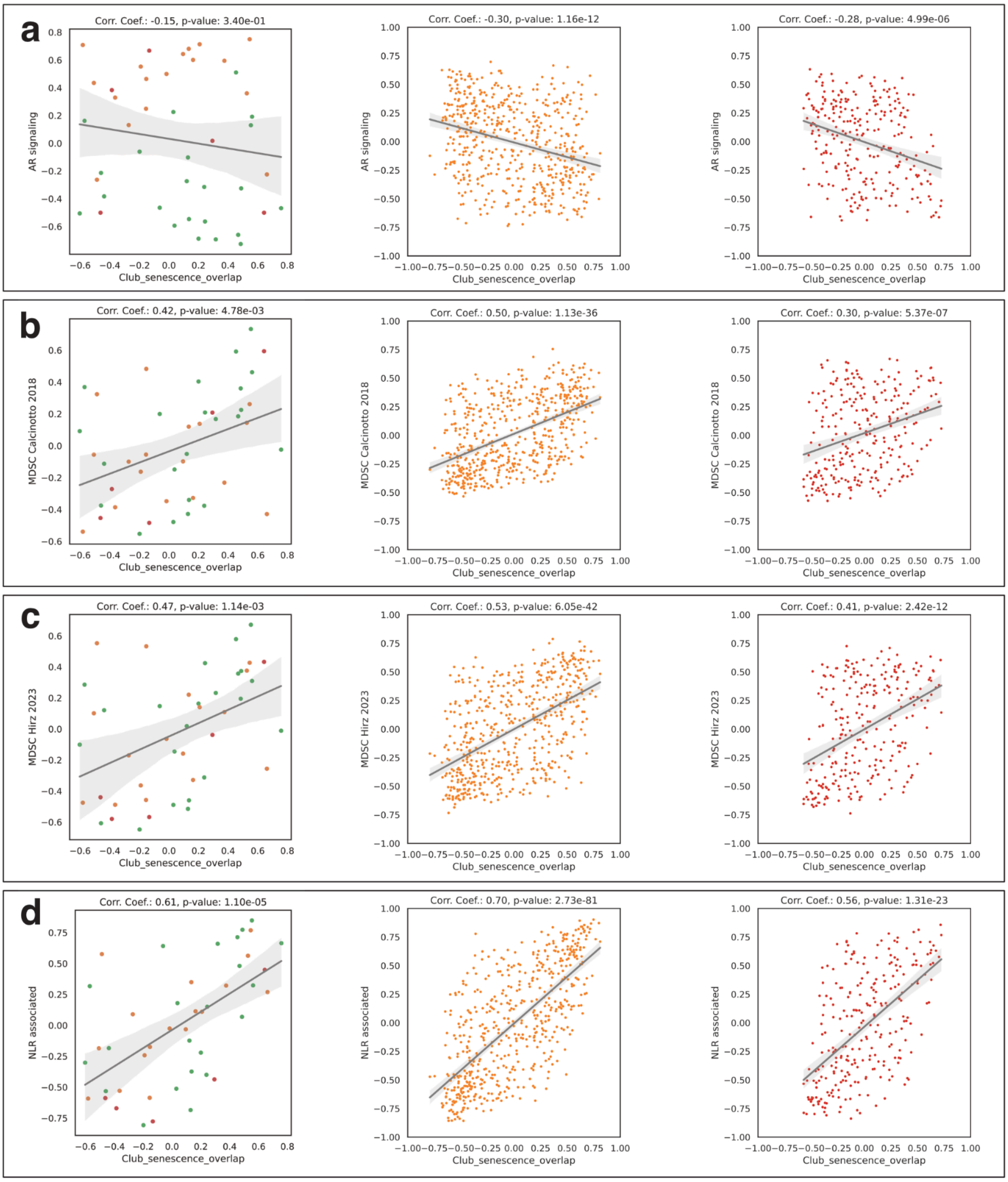


**Supplementary Figure S7.** GSVA correlations in the discovery cohort for pseudo-bulk discovery cohort ST-data (left), TCGA-PRAD (middle), and SU2C-PCF (right). Spearman correlation coefficients and their two-tailed Benjamini-Hochberg corrected p-values are displayed. **a)** AR-signaling signature[5] **b**) Calcinotto et al. 2018 MDSC signature[2] **c)** Hirz et al. 2023 MDSC signature[3] **d)** High NLR associated-signature[6]

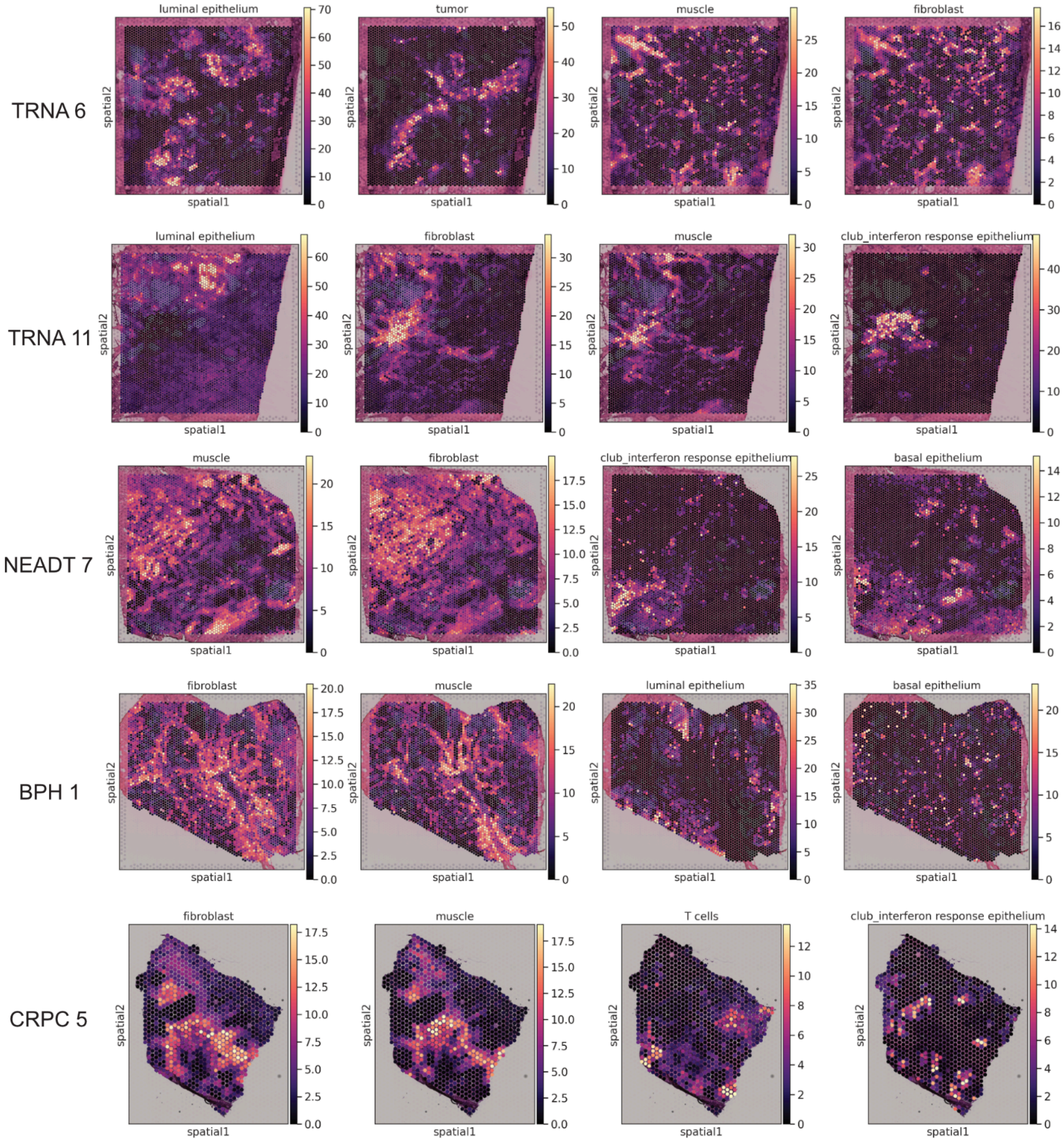


**Supplementary Figure S8.** Inferred cell count weights overlaid on spatial sections for five representative samples of the discovery cohort.

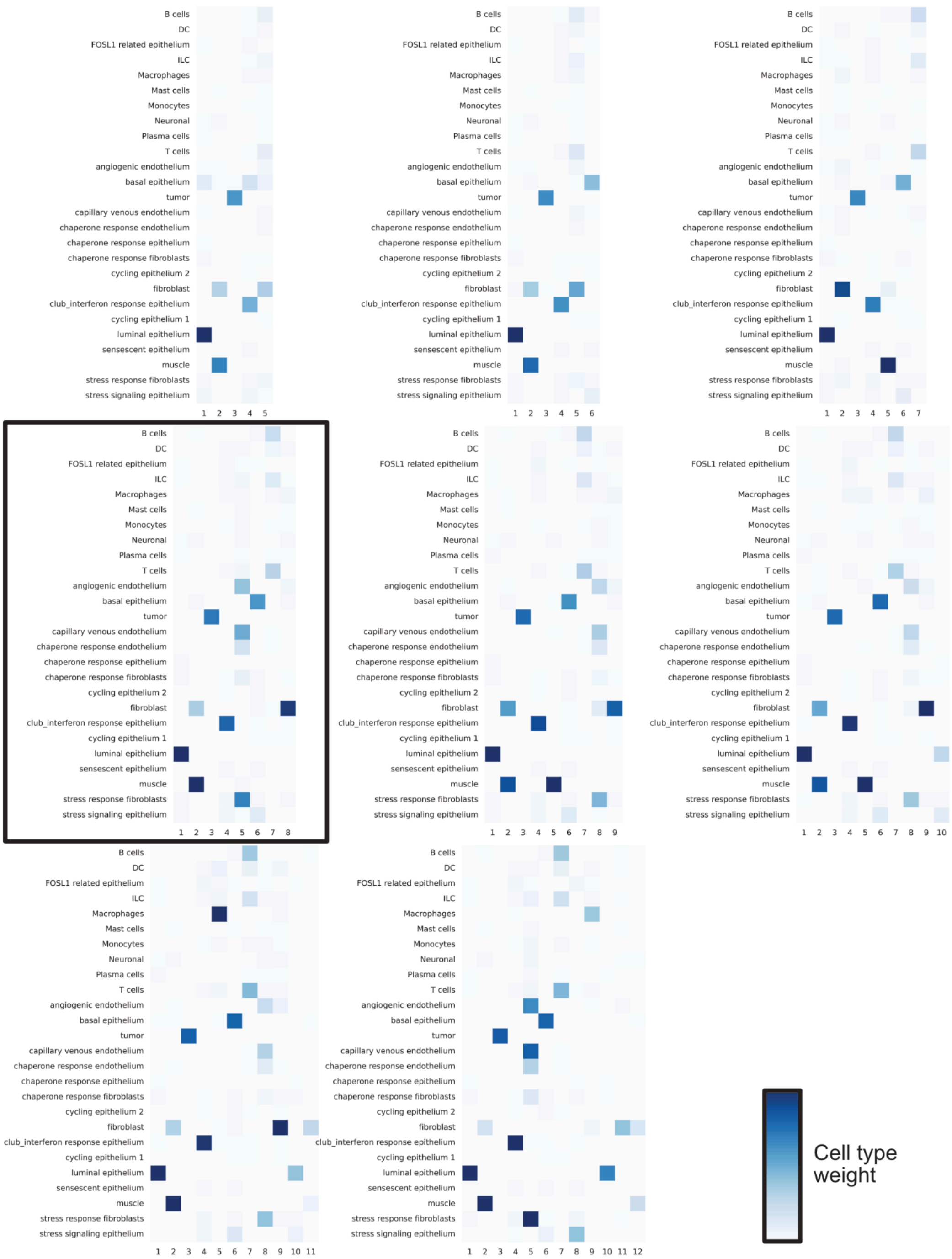


**Supplementary Figure S9.** NMF factor weights for 26 mapped cell types for the number of components k where 5 ≤ k ≤ 12. Iteration k = 8 was chosen for downstream analysis (denoted with a black rectangle). Darker blue indicates a higher contribution of a cell type to a factor.

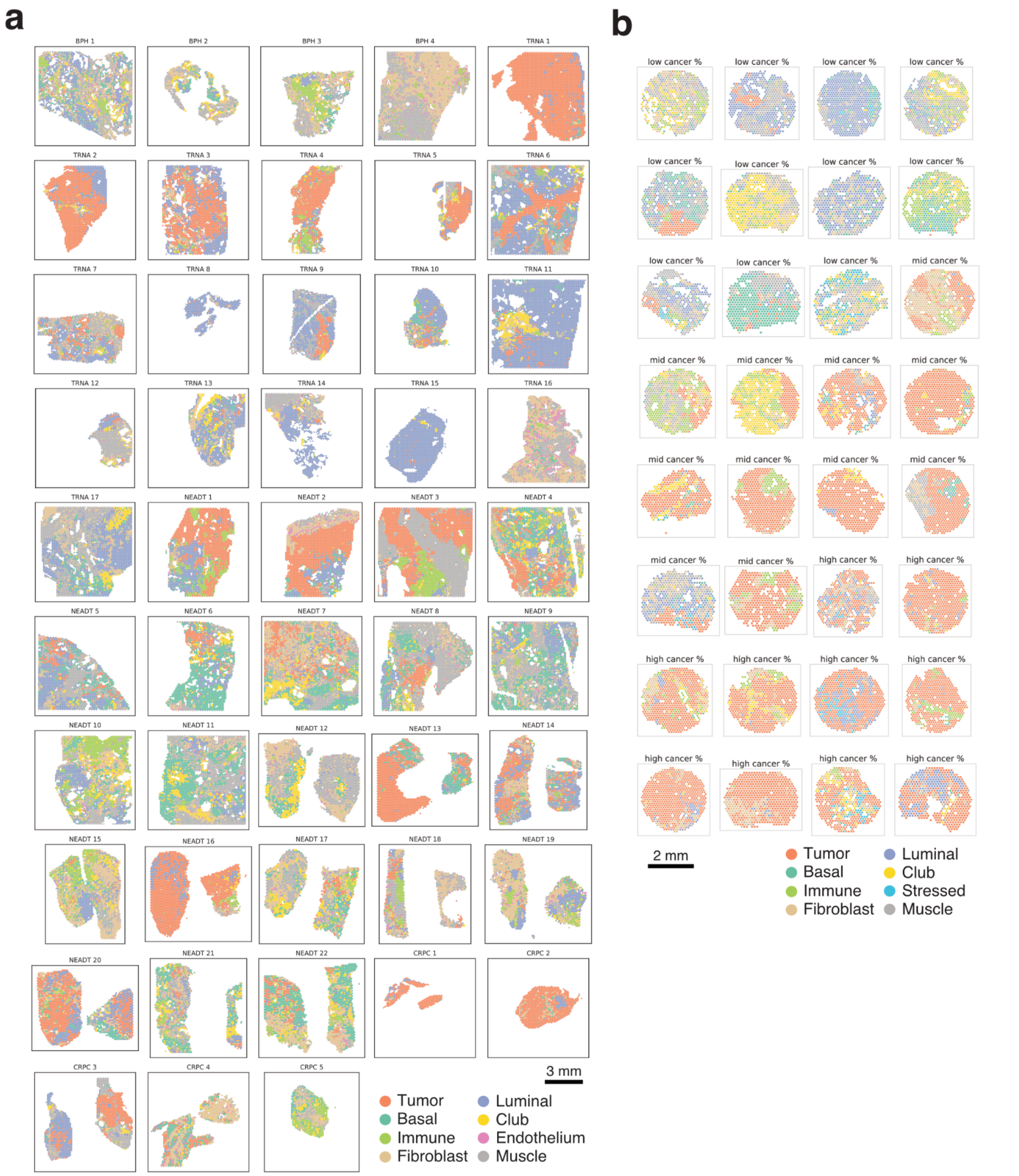


**Supplementary Figure S10.** Single-cell mapping-based (SCM) regions overlaid on the spatial sections for **a)** the discovery cohort and **b)** the validation cohort.

### Supplementary methods

#### Dataset assembly

We assembled single-cell RNA-sequencing data from 7 previously published prostate cancer studies with samples from normal prostate tissue, low- and high-grade primary tumors, as well as locally recurrent and metastatic castration-resistant prostate cancer tumors[3,7–12]. For *Dong et al. 2020, Chen et al. 2021, Song et al. 2022, Wong et al. 2022, and Hirz et al. 2023* count matrices were downloaded from the Gene Expression Omnibus (GEO) under their accession numbers (GSE137829, GSE141445, GSE176031, GSE185344, GSE181294, respectively). For Cheng et al. 2022, bam files were downloaded from the Sequence Read Archive (SRA) under accession number PRJNA699369, converted back *fastq* files using *cellranger bam2fastq* (cellranger v3.0.2), and processed using *cellranger count* (alignmed to GRCh38) to yield transcript counts. For *Chen et al. 2022*, raw FASTQ-format sequencing data were acquired from a collaborator and similarly processed identically using *cellranger count*. In total, the aggregated raw transcript counts comprised data from 64 patients, 98 samples, and 354,885 cells.

#### Dataset preprocessing

For preprocessing, normalization, and integration of the datasets, we relied on the pipeline laid out in a comprehensive single-cell RNA-seq data integration benchmark by *Luecken et al*[13]. The single-cell integration benchmark pipeline (*scib* v1.1.1) was installed as a conda environment with *scanpy* v1.9.1, python v3.7.12, and R v.4.0.5. Quality control was performed on each dataset individually. We used *scanpy* to filter cells with less than 600 UMI counts and less than 300 gene counts. Genes with less than 10 total counts across each dataset were discarded. *scanpy.pp.calculate_qc_metrics* was used to count the percentage of counts originating from mitochondrial genes, and cells with > 20% were filtered.

Scanpy's implementation of *scrublet* v0.2.3 was used to model and remove cell duplets[14]. Data normalization was performed on each dataset individually by running *scib.preprocessing.normalize* with parameters *precluster=False*, and *sparsify=False*. Following normalization, the datasets were concatenated into a single object with *anndata.concat* v0.8.0 using the default parameters. The resulting aggregate dataset consisted of 327,771 cells and 14,819 genes.

#### Dataset integration

*scib.preprocessing.scale_batch* was next used to jointly scale the data with the dataset of origin used as the batch key. Highly variable genes (HVGs) were identified by running *scib.preprocessing.hvg_batch* with parameters *target_genes=2000*, *flavor='seurat',* and the dataset of origin as batch key. To perform dataset integration, we used the *scvi* method in *scvi-tools* v0.1.16.1. We used the unnormalized counts of the identified HVGs and dataset as batch keys. A model was constructed using *scvi.model.SCVI* with the following parameters: *n_layers=2*, *n_latent=30*, *gene_likelihood='nb'*. The model was trained on an NVIDIA Tesla V100 architecture GPU. A 30-dimensional latent representation of the dataset was extracted from the trained model. This representation was then graph-clustered by using *scanpy.pp.neighbors* and *scanpy.tl.leiden* with the default parameters. A UMAP representation was constructed using *sc.tl.umap*.

#### Broad cell-type annotation

Leiden clustering of the 30-dimensional latent space resulted in 50 unique clusters (VI_clusters, **Supplementary Figure S11**). From these clusters, we discarded those that had more than 80% of their cells originating from a single sample. This resulted in the exclusion of 16 clusters (42,917 cells), or 41.8%, 11.6%, and 5.3% of cells originating from CRPC, PCa, and normal samples, respectively.

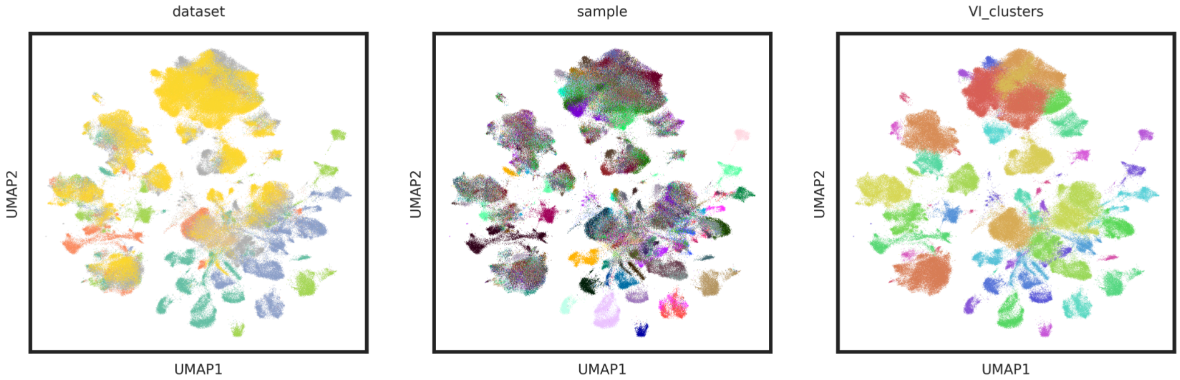


***Supplementary Figure 11.*** *UMAP representation of the 30-dimensional latent space inferred through integration with scvi. Individual cells are colored by dataset (7), sample (98), and cell clusters yielded by Leiden clustering of the latent space (50).*

To annotate these data, we assembled a set of consensus gene markers for 11 cell types from the original publications used in our integrated dataset. This categorization was intentionally broad and contained epithelial, endothelial, fibroblast, muscle, and neuronal cell type markers. The immune cell markers were for mast, T-, B-, myeloid-, dendritic-, and plasma cells.

Cell types were annotated using the following procedure. First, a dendrogram relating the 34 remaining clusters to each other was calculated from the 30-dimensional latent space with *scanpy.tl.dendrogram*. The gene expression was then plotted using *sc.pl.dotplot*, with the cell type consensus gene markers on one axis and clusters ordered according to the dendrogram on the other (**Supplementary Figure S12**). Starting from left to right, each cluster was then annotated as one cell type with ambiguous cases being annotated according to their neighbors on the dendrogram. This procedure resulted in 10 unique cell-type clusters, where fibroblast and muscle categories were merged into one due to their similarity in terms of marker gene expression.

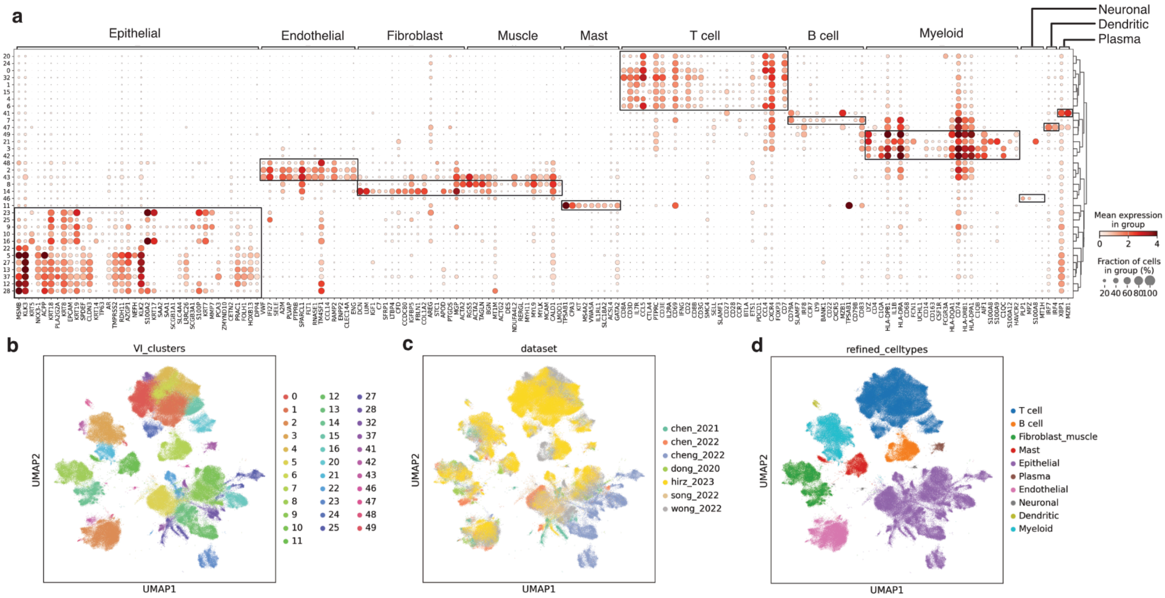


***Supplementary Figure S12.** Cell type identification of integration-derived VI_clusters. **a)** Expression of canonical marker genes in each cluster. Cell types are shown on top, individual genes on the bottom. Rectangles drawn on each cluster at specific cell types indicate inferred cluster identities. **b,c,d)** 34 post-exclusion VI_clusters, datasets, and broad cell-type annotations are shown in the UMAP representation of the 30-dimensional latent space.*

#### Annotating immune cell identity with celltypist

We used *celltypist* v1.6.0, a machine-learning model optimized for immune cell identity annotation of tumor-derived single-cell RNA-seq data[15]. The raw counts of cells annotated as T-, B-, mast, plasma, dendritic, and myeloid cells were jointly normalized and log-transformed using *scanpy*, after which they were annotated using *celltypist.annotate* with *Immune_All_High* used as the model. This mode examined the expression of 6639 genes and compared them to a high-hierarchy immune cell annotation derived from 20 tissues across 18 studies. The model was trained using thresholds 0.99 for probability, and 100 for the number of cells. The *celltypist* predictions corroborated T-, B-, mast-, plasma- and dendritic cell annotations given in the upstream analysis, while also introducing innate lymphoid cells (ILCs), macrophages, and monocytes, resulting in 98,662 annotated immune cells across these eight categories (**Supplementary Figure S13**).

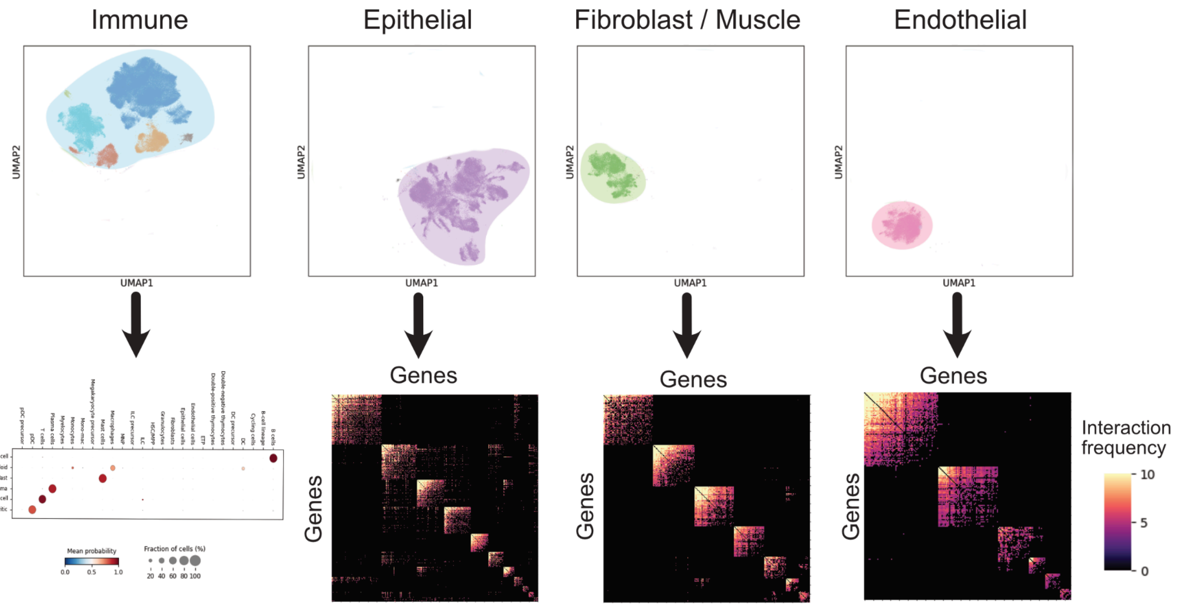


***Supplementary Figure S13.*** *NMF-based annotation of broadly categorized cell types. The broad annotations were divided into immune, epithelial, fibroblast/muscle, and endothelial subsets. The immune subset underwent annotation using celltypist, while the other 3 underwent an NMF-based algorithmic annotation.*

#### Using non-negative matrix factorization to infer cell-type specific intra-sample heterogeneity

Based on the broadly annotated cell types, we set out to find the shared intra-tumor heterogeneity using a previously described non-negative matrix factorization (NMF) approach[16]. We ran the following algorithm for the epithelial, fibroblast/muscle, and endothelial cell types: For each sample individually, cells labeled as the cell type of interest were subsetted. Using the unnormalized gene counts from these cells, genes with less than 10 counts were removed using *scanpy*. These counts were then normalized, scaled, and subsetted to HVGs using *scanpy.pp.normalize_total*, *scanpy.pp.scale*, and *scanpy.pp.highly_variable_genes*, respectively. HVGs were determined using parameters *n_top_genes=2000, flavor='seurat_v3', subset=True*, and *layer=counts*. Negative scaled expression values were set to 0. On these data, *nimfa* v1.4.0 was used to perform non-smooth NMF for a decreasing number of components *k* ranging from 25 to 5. Samples with less than 100 cells annotated as the cell type in question were discarded (epithelial: 12, fibroblast/muscle: 52, endothelial: 44 samples discarded). After each iteration, the integrity of the resulting components was checked by observing their unique genes as follows.

One factor at a time, genes were ranked in decreasing order according to the weight associated with this factor. This weight was then compared across all the other factor weights associated with this gene. If the factor of interest had the highest weight for this gene, the gene was annotated as specific to the factor of interest. Iterating through the gene list, this procedure was repeated until a factor other than the factor of interest had a larger weight for the gene in question, resulting in a set of genes specific to the factor of interest. By repeating this process across all factors, a set of factor-specific genes was attached to each factor. If there were less than 5 genes in any of the factor-specific gene sets, NMF was performed using a smaller number of components by setting *k=k-1*. The final result of the algorithm was a set of genes for *k* components for each individual sample.

Following the NMF algorithm, redundant factors were discarded by observing the gene set overlap across factors (samples, as each gene was only present in one factor per sample). Factors that had a minimum Jaccard index of 0.05 with 2 or more other factors were retained. These factors were then used to construct a gene-interaction matrix, where each entry represented the number of factors in which the two genes co-occurred.

#### Defining gene modules and their activity in single-cell data

The gene-interaction matrix could be interpreted as a graph, where each gene is a node and each entry, representing the number of co-occurrences in the NMF factors, is an edge connecting two nodes. Gene-gene connections that were observed in less than 4 factors and genes with less than 5 unique connections were determined redundant and removed. Next, *igraph* v0.10.3 was used to build the gene adjacency matrix into an undirected graph, which was then clustered with *igraph.community_leiden*. Clusters with 10 or more genes were included as the final gene modules (**Supplementary Table S2**). To annotate cells according to these gene modules, *scanpy.tl.score_genes* was run for each module in each cell that belonged to the corresponding broad cell type annotation. Each cell was given a label according to its highest-scoring module. Modules with less than 1000 top-scoring cells were omitted.

In total, this approach yielded 124,703 annotated cells across ten epithelial, four fibroblast/muscle, and three endothelial subtypes. Combined with the annotated immune cells and neuronal cells, this intricately annotated single-cell reference comprised 223,881 cells across 26 cell states. These data were used as a single-cell mapping reference for deconvolving the spatial transcriptomics data, as described in **Methods**.
